# MYCN inhibits TrkC-mediated differentiation in neuroblastoma cells via disruption of the PKA signalling pathway

**DOI:** 10.1101/2024.08.07.606961

**Authors:** Stephanie Maher, Kieran Wynne, Vadim Zhernovkov, Melinda Halasz

**Affiliations:** Systems Biology Ireland, School of Medicine, University College Dublin, Belfield, Dublin, Ireland; Conway Institute of Biomolecular and Biomedical Research, University College Dublin, Belfield, Ireland

## Abstract

Neuroblastoma is a complex paediatric cancer with a spectrum of clinical outcomes ranging from spontaneous regression to aggressive metastatic disease. Low-risk patients achieve over 90% survival with no or minimal treatment, while high-risk patients face less than 50% survival despite intensive multimodal therapy. Half of the high-risk cases harbour amplification of the *MYCN* oncogene. In addition to MYCN status, Trk receptors have also been linked to prognosis. TrkA expression is seen with low-risk cases while TrkB expression often occurs in high-risk MYCN-amplified NB. While TrkA and TrkB are well studied in NB, the role of TrkC in neuroblastoma genesis is not clear. Therefore, this study investigates the interplay between MYCN status and NT-3/TrkC signalling in neuroblastoma. Using a panel of neuroblastoma cell lines with varying MYCN levels, we found that TrkC activation leads to neuronal differentiation of MYCN non-amplified cells, whereas it promotes proliferation of MYCN-amplified cells. Temporal phosphoproteomics revealed differential activation of the PKA pathway, which was crucial for TrkC-mediated differentiation. Manipulating the PKA pathway altered cell fate outcomes, underscoring its role. In MYCN-amplified cells, MYCN knockdown increased PKA and CREB activity, shifting the phenotype towards differentiation. Analysis of neuroblastoma patient data showed lower expression of PKA pathway genes in MYCN-amplified tumours. Additionally, miR-221, upregulated by MYCN, was identified as a suppressor of the PKA/CREB pathway. These findings highlight the context-dependent nature of NT-3/TrkC signalling influenced by MYCN; and suggest therapeutic potential in targeting the PKA pathway to induce differentiation of high-risk MYCN-amplified neuroblastoma.

## Introduction

Neuroblastoma is a paediatric cancer originating from neural crest cells of the developing sympathetic nervous system. Extreme clinical heterogeneity is a hallmark of neuroblastoma, ranging from spontaneous tumour regression to aggressive metastatic progression (1,2). Patients are stratified into low-, intermediate- and high-risk groups defined by prognostic factors, including age at diagnosis, tumour stage, histological features, and cytogenetic findings (3,4). Patients with low-risk disease typically receive clinical observation or surgical resection of the primary tumour, achieving a 5-year overall survival rate exceeding 90%. Some of these tumours may spontaneously regress through apoptosis or neuroblast differentiation.(5). Conversely, approximately 50% of patients are considered high-risk and harbour aggressive, metastatic neuroblastomas that often resist treatment and subsequently relapse. Despite intensive multimodal therapeutic approaches, the 5-year overall survival rates for these high-risk patients remain below 50% (3,5).

The neurotrophic tyrosine receptor kinase (NTRK) family, comprising TrkA, TrkB and TrkC as well as their respective ligands NGF, BDNF and NT-3, are known to be differentially expressed in the divergent clinical phenotypes we observe in neuroblastoma patients(6–9). TrkA is associated with low-stage disease and demonstrates high expression in the spontaneous regression phenotype (6,10,11). Conversely, TrkB is associated with aggressive high-risk neuroblastomas, actively promoting angiogenesis, proliferation, and invasion (12–15).

TrkC signalling has also been investigated in the context of neuroblastoma albeit to a lesser extent than TrkA and TrkB. Previous research reports high expression of TrkC in low stage neuroblastomas that have a favourable outcome and an association with TrkA expression (9,16). Additional studies also allude to its potential oncogenic abilities; a subset of advanced stage 4 neuroblastomas have been identified with high levels of both NT-3 and TrkC. This co-expression is thought to contribute to an autocrine mechanism, driving the survival and proliferation of neuroblastoma cells (17). Beyond neuroblastoma, the role of TrkC in cancer is multifaceted. In some cancers, such as medulloblastoma and colon cancer, TrkC expression is associated with favourable outcomes, suggesting a potential tumour-suppressive function. However, the situation is reversed in breast cancer and leukaemia where TrkC exhibits oncogenic signalling. This duality underscores the context-dependent nature of NT-3/TrkC signalling activity, indicating that the effects of TrkC on cell behaviour are highly influenced by the cellular and molecular environment in which it operates (18–22).

Integral to the tumorigenesis of neuroblastoma is the MYCN oncogene. During normal embryogenesis, MYCN is expressed transiently in the ventral-lateral migrating cells of the neural crest that will become the sympathetic ganglia and mediates cellular proliferation, migration and differentiation through the activation and repression of genetic targets (23,24). Amplification of MYCN occurs in approximately 20-25% of neuroblastomas and independently classifies a high-risk diagnosis. These tumours show a highly aggressive phenotype and compared to MYCN non-amplified neuroblastoma, a considerably worse outcome (25–27). In addition to promoting proliferation, MYCN amplification promotes angiogenesis, tumour metastasis and evasion of immune surveillance (24). Amplification also potently inhibits differentiation pathways and maintains self-renewal and pluripotency of cells, creating a highly aggressive tumour environment. Not surprisingly, MYCN amplification is found in 30-40% of stage 3 and 4 neuroblastoma patients (28).

Currently, there is limited understanding of the interplay between MYCN status and NT-3/TrkC signalling in neuroblastoma. The dual nature of TrkC’s effects, acting as both an oncogene and a tumour suppressor depending on the cellular context, highlights the need for precise molecular characterisation of its signalling. This study seeks to elucidate the dynamics of the NT-3/TrkC signalling network and its impact on cell fate decisions in neuroblastoma, with a focus on how the oncogenic driver MYCN influences this signalling landscape.

## Results

### NT-3/TrkC signalling directs different cell fates in neuroblastoma cells with differential MYCN levels

To study TrkC receptor signalling in neuroblastoma, we first established a panel of neuroblastoma cell lines that stably express the NTRK3 gene, encoding the TrkC protein. We chose the SH-SY5Y, NBLS and NLF neuroblastoma cell lines (Fig.1A, Table 1.) to transfect with the NTRK3 gene on the basis that they do not exhibit basal TrkC expression (Fig.1B), and the MYCN status of the cells is different. SH-SY5Y cells are MYCN non-amplified, NBLS cells overexpress MYCN from a single gene copy, and NLF cells harbour MYCN amplification. In addition, the MYCN levels are comparable in NBLS and NLF cells (Fig.1A). MYCN is the main oncogenic driver in neuroblastoma and MYCN amplification status is independently prognostic of a high-risk disease. By using a panel of neuroblastoma cell lines with varying MYCN levels we can therefore assess the influence of MYCN status on the TrkC receptor signalling network in neuroblastoma.

**Figure 1.**
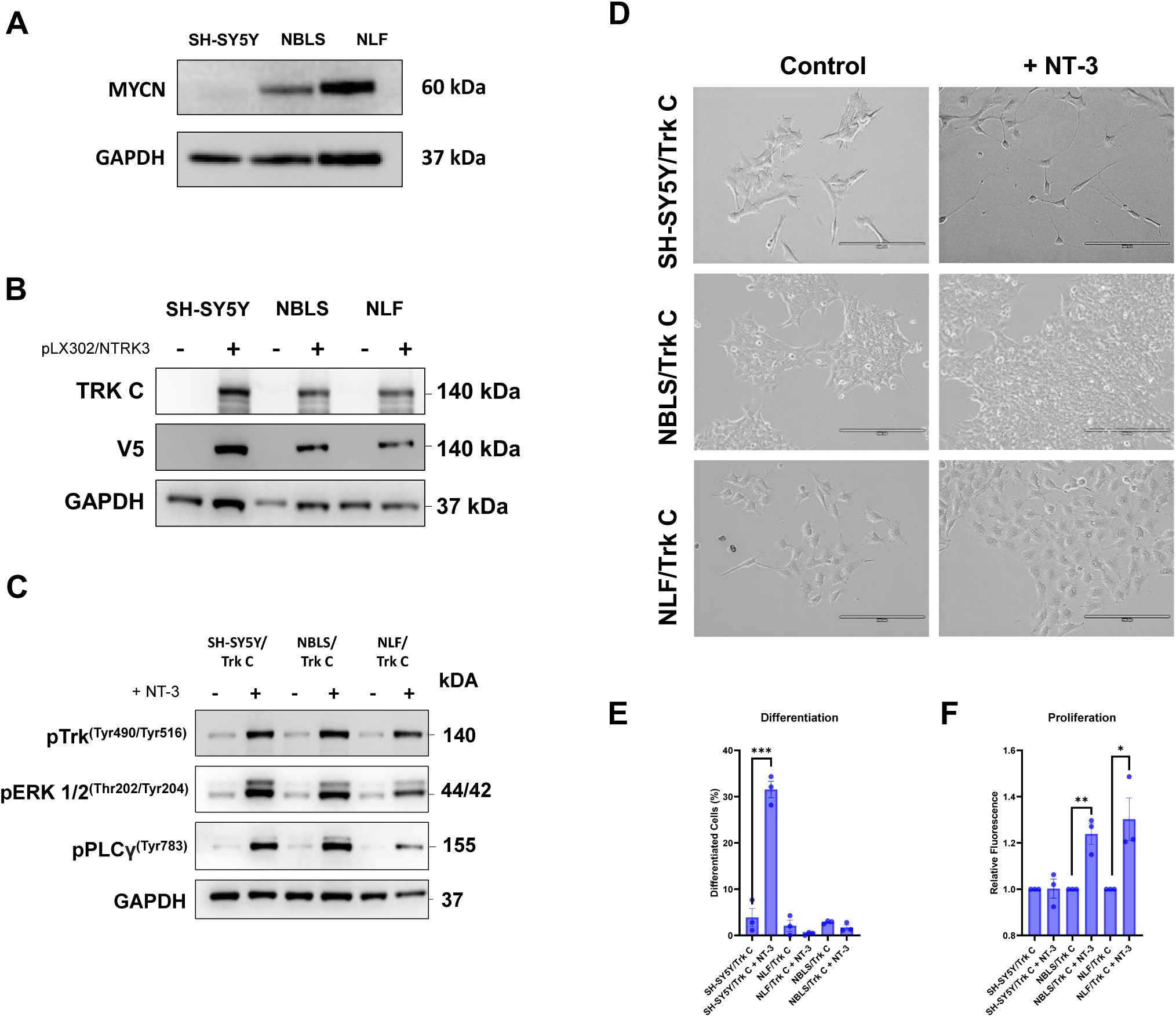
TrkC drives divergent cellular phenotypes in neuroblastoma cells with different MYCN status. **A)** MYCN expression of SH-SY5Y, NBLS and NLF parental neuroblastoma cell lines by Western blotting. **B)** TrkC and V5-tag expression in parental neuroblastoma cell lines and pLX302/NTRK3 transfected cells. **C)** Western blot analysis of phospho-TrkA ^(Tyr490)^/TrkB ^(Tyr516)^, phospho-ERK ^(Thr202/Tyr204),^ phospho-PLCγ^(Tyr783)^ following stimulation of SH-SY5Y/TrkC, NBLS/TrkC, NLF/ TrkC cells with 100 ng/ml of NT-3 for 10 mins. GAPDH acted as loading control for Western blot experiments. **D)** Representative images of phenotypic observation by light microscopy 5 days post NT-3 (100 ng/ml) stimulation in SH-SY5Y /TrkC, NLF/TrkC and NBLS/TrkC neuroblastoma cells. Magnification: X20; scalebar: 200 µm. **E)** Quantification of neuronal differentiation following NT-3 (100 ng/ml) treatment for 5 days. **F)** Quantitation of mean fluorescence as a measure of cell proliferation at 72h post NT-3 (100 ng/ml) stimulation using CyQuant assay (n =3). Data shown as mean +/-SD, (p < 0.05 = *, p < 0.001 = **, p < 0.0001 = ***).

**Table 1.** Genetic profile of neuroblastoma cell lines used in the project. (**41**).

| Cell Line | MYCN status | 1p36 del | 3p26 del | 11q23 del | 17q unbalanced gain | ALK mutation | p53 mutation |
| --- | --- | --- | --- | --- | --- | --- | --- |
| SH-SY5Y | Non-amplified | – | – | + | + | F1174L | WT |
| NBL-S | Non-amplified, MYCN overexpression | – | – | + | + | WT | WT |
| NLF | Amplified | + | + | + | + | WT | V203M |
| IMR-32 | Amplified | + | + | + | + | WT | WT |

Activation of the TrkC receptor with its cognate ligand, NT-3 resulted in phosphorylation of the receptor and activation of downstream signalling pathways including ERK and PLCy (Fig 1C). This is a putative consequence of Trk receptor activation and is in line with previously published research, thus deeming our cell system suitable to study TrkC receptor signalling in neuroblastoma (29–31).

To assess cell fate decisions regulated by TrkC signalling, we assessed neuronal differentiation and proliferation upon NT-3 treatment. These are the two predominant cell fates in neuroblast development correlating clinically with spontaneous regression and aggressive metastatic progression, respectively. Cell fate decisions directed by NT-3/TrkC signalling were different in the three cell lines. SH-SY5Y/TrkC cells stimulated with NT-3 resulted in neuronal differentiation and no significant change in cell number compared with untreated control (Fig 1D-F). In contrast, NBLS/TrkC and NLF/TrkC cells with MYCN overexpression and MYCN amplification, respectively, did not undergo differentiation when treated with NT-3. Likewise, NBLS/TrkC and NLF/TrkC cells also displayed a significant increase in cell proliferation. This characterisation highlights a difference in cell fate decisions mediated by NT-3/TrkC across neuroblastoma cell lines, and the different cell fates observed might be due to the different MYCN levels of the cells.

### Temporal phosphoproteomics of NT-3/TrkC signalling reveals differential PKA pathway activation in the neuroblastoma cells with different cell fates

To understand the dynamic changes in the downstream NT-3/TrkC signalling network that may be contributing to the divergent cell fate decisions observed, a quantitative phosphoproteomics approach was used. Temporal profiling of SH-SY5Y/TrkC, NBLS/TrkC and NLF/TRKC cells was carried out by stimulating the cells with NT-3 for 0, 10, 45 minutes and 24 hours (Fig. 2A). This range of timepoints facilitated the capture of early, intermediate, and late phosphorylation events.

**Figure 2.**
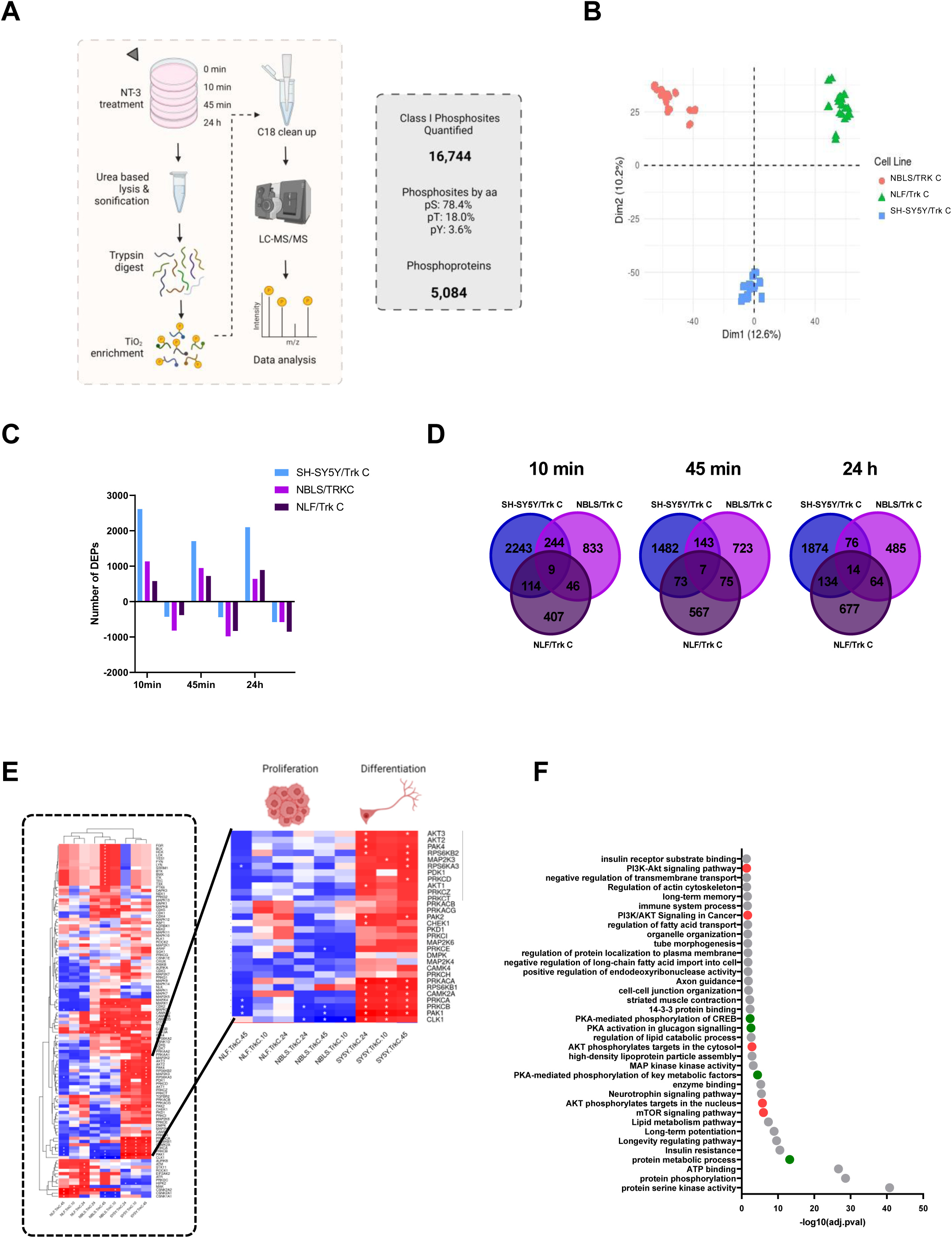
Phosphoproteomics analysis reveals differences in NT-3/TrkC signalling across SH-SY5Y/TrkC, NBLS/TrkC and NLF/TrkC cells. **A)** Quantitative phosphoproteomics experimental workflow for the analysis of NT-3/TrkC signalling in neuroblastoma cells and summary of phosphoproteome data with information on number of identified phosphorylation sites and phosphoproteins (3 cell lines, 4 time points, 3 biological and 2 technical replicates resulting in 72 MS runs). **B)** Principal component analysis (PCA) plot of all phosphoproteomics samples (n = 72) **C)** Number of upregulated and downregulated differently expressed phosphosites (DEPs) after 10’, 45’ or 24h NT-3 treatment (adjusted p-value < 0.05 and absolute fold change > 1.5). **D)** Venn diagram of overlapping DEPs between each cell line at 10’, 45’ and 24h NT-3 treatment. **E)** Kinase substrate enrichment analysis with KSEAapp of DEPs in all samples FDR < 0.05, m.cutoff = 3. **F)** Pathway enrichment analysis for all upregulated DEPs in SH-SY5Y/TrkC cells, p-value < 0.05 (70)

We identified and quantified 25,098 phosphorylated sites, 16,744 of which were confidently localized to serine (78.4% of the total), threonine (18.0%), or tyrosine (3.6%) residue in the peptide sequence class I with high reproducibility between replicates. Principle Component Analysis (PCA) of all phosphoproteomics samples resulted in distinct clustering based on the parental cell line (Fig 2B). Furthermore, assessment of differently expressed phosphosites (DEPs) revealed that SH-SY5Y/TrkC cells exhibit the highest number of upregulated DEPs compared to the other cell lines at each timepoint. NBLS/TrkC cells demonstrated the highest number of significantly downregulated phosphosites at 10 and 45 minutes of NT-3 treatment, while NLF/TrkC cells had the highest number of downregulated phosphosites after 24 hours of NT-3 stimulation (Fig 2C). Moreover, SH-SY5Y/TrkC cells also displayed the greatest number of unique differentially expressed phosphosites at each timepoint of NT-3 treatment when comparing the overlapping DEPs (Fig.2D). Taken together, the trends in the phosphoproteomics data correspond with the observed cell fate decisions. SH-SY5Y/TrkC cells demonstrate a strong positive response to NT-3 stimulation and harbour a distinctive phosphoproteome compared to NBLS/TrkC and NLF/TrkC cells. This enhanced response may be attributed to the known role of MYCN as a global suppressor of cell signalling in neuroblastoma, resulting in predominantly upregulated pathways in the MYCN non-amplified SY5Y/TrkC cells compared to downregulated pathways in the MYCN overexpressing NBLS/TrkC and MYCN-amplified NLF/TrkC cells.

As phosphoproteomics datasets can be large and complex to delineate, kinase substrate enrichment analysis (KSEA) was employed to focus on the most relevant phosphorylation changes in the NT-3/TrkC phosphoproteome data by linking the phosphorylation changes to key regulatory kinases, thus reducing data complexity.

Firstly, KSEA analysis showed kinases clustering on the basis of TrkC cells that differentiate (SH-SY5Y/TrkC) or proliferate (NBLS/TrkC and NLF/TrkC) when stimulated with NT-3 suggesting the downstream signalling network to be mediating the different cell fate decisions. Protein Kinase A (PKA) and Protein Kinase B (AKT) were identified as key regulators of SH-SY5Y/TrkC signalling that were not enriched in NBLS/TrkC and NLF/TrkC cells (Fig. 2E).

Furthermore, pathway enrichment analysis of phosphoprotein kinases upregulated in SH-SY5Y/TrkC cells showed several expected pathways including “neurotrophin signalling pathway”, “protein phosphorylation” and “protein serine kinase activity”. It also showed enrichment for processes relating to neuronal differentiation including “axon guidance”, “regulation of actin cytoskeleton” and “long term potentiation”. There were also several pathways relating to metabolism including “regulation of fatty acid transport”, “regulation of lipid catabolic process”, “protein metabolic process”, “lipid metabolism pathway”. Most notably, there was an enrichment for terms relating to the PKA and AKT pathways including “PI3K-Akt signalling pathway”, “AKT phosphorylation targets in the cytosol”, “mTOR signalling pathway” and “PKA-mediated phosphorylation of CREB”, “PKA activation in glucagon signalling”, “PKA-mediated phosphorylation of key metabolic factors”, further suggesting these pathways to play a role in NT-3/TrkC mediated neuronal differentiation of SH-SY5Y/TrkC cells (Fig 2F).

### Manipulation of the PKA pathway alters TrkC-mediated cell fate decisions in neuroblastoma

To test if the differential PKA signalling drives the different cell fate decisions observed, we inhibited or activated PKA signalling in SH-SY5Y/TrkC or NBLS/TrkC and NLF/TrkC cells, respectively. Pharmacological inhibition of PKA signalling by H89 HCl resulted in increased cell number and attenuation of neuronal differentiation in NT-3 treated SH-SY5Y/TrkC cells (Fig 3A-B,D). Conversely, activation of the PKA pathway in NT-3 treated NBLS/TrkC and NLF/TrkC cells by dibutyryl cyclic AMP (db-cAMP, a cell-permeable cAMP analogue), resulted in neuronal differentiation and inhibition of cell proliferation (Fig. 3A, 3C-D-,3F-G), highlighting a role for the PKA pathway in steering TrkC mediated neuroblast cell fate decisions.

**Figure 3.**
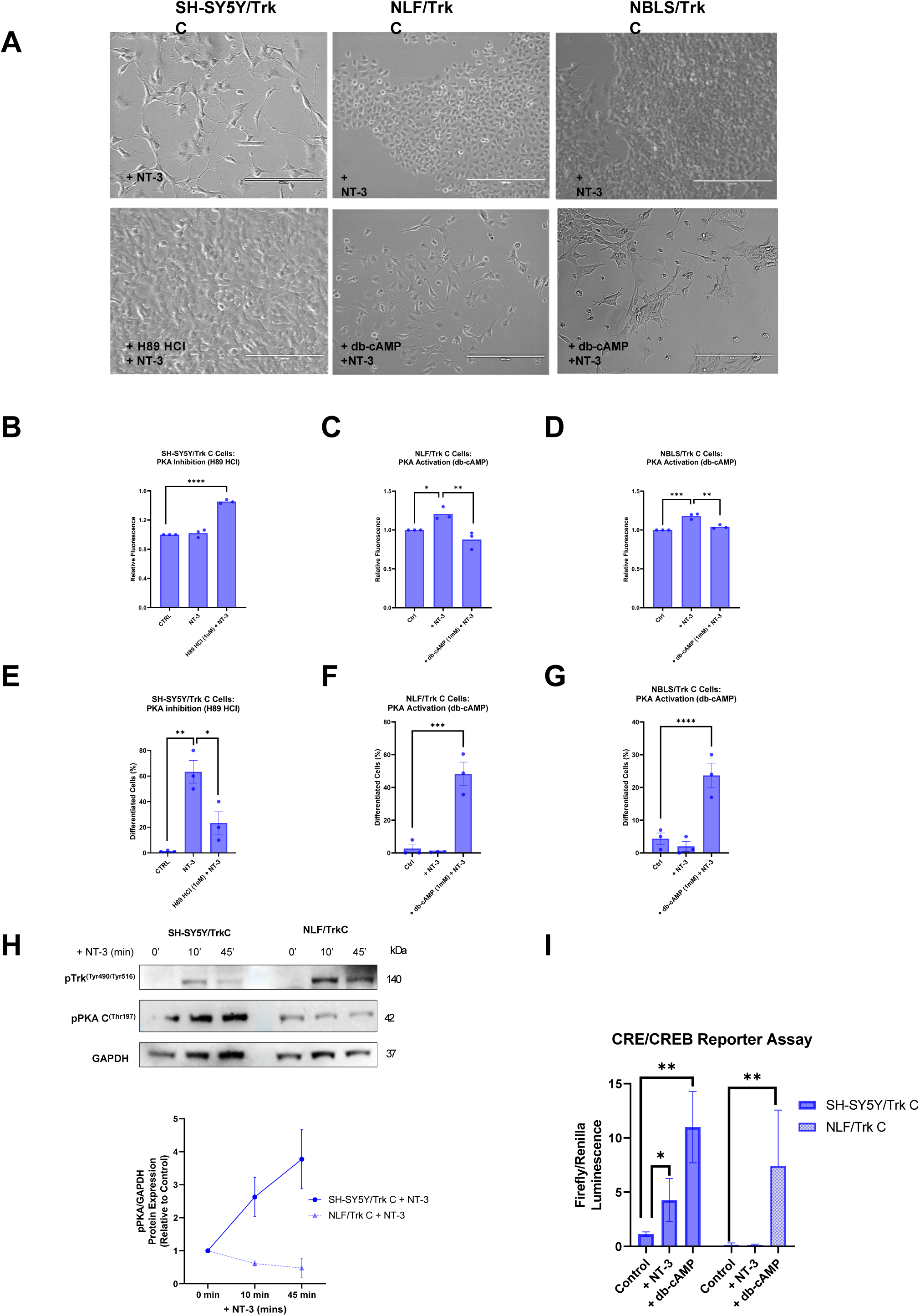
Manipulation of the PKA pathway alters TrkC-mediated cell fate decisions in neuroblastoma cells. **A)** Representative images of cells following 5-day treatment with NT-3 (100 ng/ml) and/or H89 HCl (1 μM), db-cAMP (1 mM) in SH-SY5Y/TrkC, NLF/TrkC and NBLS/TrkC cells. Magnification: X20; scalebar: 200 µm. **B)** Measure of cell proliferation using CyQuant assay following 72 h of treatment as in A) in SH-SY5Y/TrkC, **C)** NLF/TrkC and **D)** NBLS/TrkC cells. Data was quantified as mean fluorescence as a measure of relative cell number. **E)** Percentage of differentiated neurons following 5 days of treatment as in A) in SH-SY5Y/TrkC, **F)** NLF/TrkC and **G)** NBLS/TrkC cells. **H)** Western blot analysis of phospho-TrkA ^(Tyr490)^/TrkB ^(Tyr516)^ and phospho-PKA C^(Thr197)^ in SH-SY5Y/TrkC and NLF/TrkC cells following 0-10-45 min treatment with NT-3 (100 ng/ml). GAPDH acted as loading control. **I)** Measure of CRE/CREB transcriptional activity by luciferase reporter assay following NT-3 stimulation (100 ng/ml) in SH-SY5Y/TrkC and NLF/TrkC cells, db-cAMP (1 mM) was used as positive control. Data is presented as mean +/-SD of at least three independent experiments (n=3) (p < 0.05 = *, p < 0.001 = **, p < 0.0001 = ***).

In its inactive state, PKA exists as a tetrameric holoenzyme composed of two catalytic (C) subunits (three isoforms: C-α, C-β, C-γ) and two regulatory (R) subunits (four isoforms: RI-α, RI-β, RII-α, RII-β). Activation results in a conformational change that cause the release and activation of catalytic subunits from the regulatory subunits. On assessment of signalling, PKA C became phosphorylated at Thr197 upon NT-3 stimulation in SH-SY5Y/TrkC cells. In contrast, PKA C demonstrated no significant change in Thr197 phospho-activity in NLF/TrkC cells when stimulated with NT-3 (Fig. 3H).

A putative event in PKA C downstream signalling is activation of the cAMP response element-binding protein (CREB), and initiation the transcription of target genes containing cAMP response elements (CREs) in their promoter regions. Evaluation of CRE/CREB transcriptional factor activity by luciferase reporter assay showed an increase in CRE/CREB transcriptional activity upon NT-3 stimulation in SH-SY5Y/TrkC cells whereas no increase in NLF/TrkC cells compared to untreated control confirming differences in the regulation of this pathway across cell lines (Fig. 3I).

### MYCN status influences TrkC downstream signalling and cell fate decisions in neuroblastoma cells

To ascertain whether differences in cell fate decisions and activation of the cAMP/PKA/CREB pathway between SH-SY5Y/TrkC cells and NLF/TrkC cells was primarily attributed to the difference in MYCN status rather than other genomic characteristics, siRNA-mediated knockdown (KD) of MYCN was employed. Following 24 hours, MYCN protein expression levels were reduced by ∼60% in NLF/TrkC cells (Fig. 4A). Consequently, this reduction in MYCN levels also resulted in increased expression of total PKA C-α and CREB in NLF/TrkC cells (p = 0.0481 & 0.0293, respectively) suggesting an inverse relationship between these proteins (Fig. 4A).

**Figure 4.**
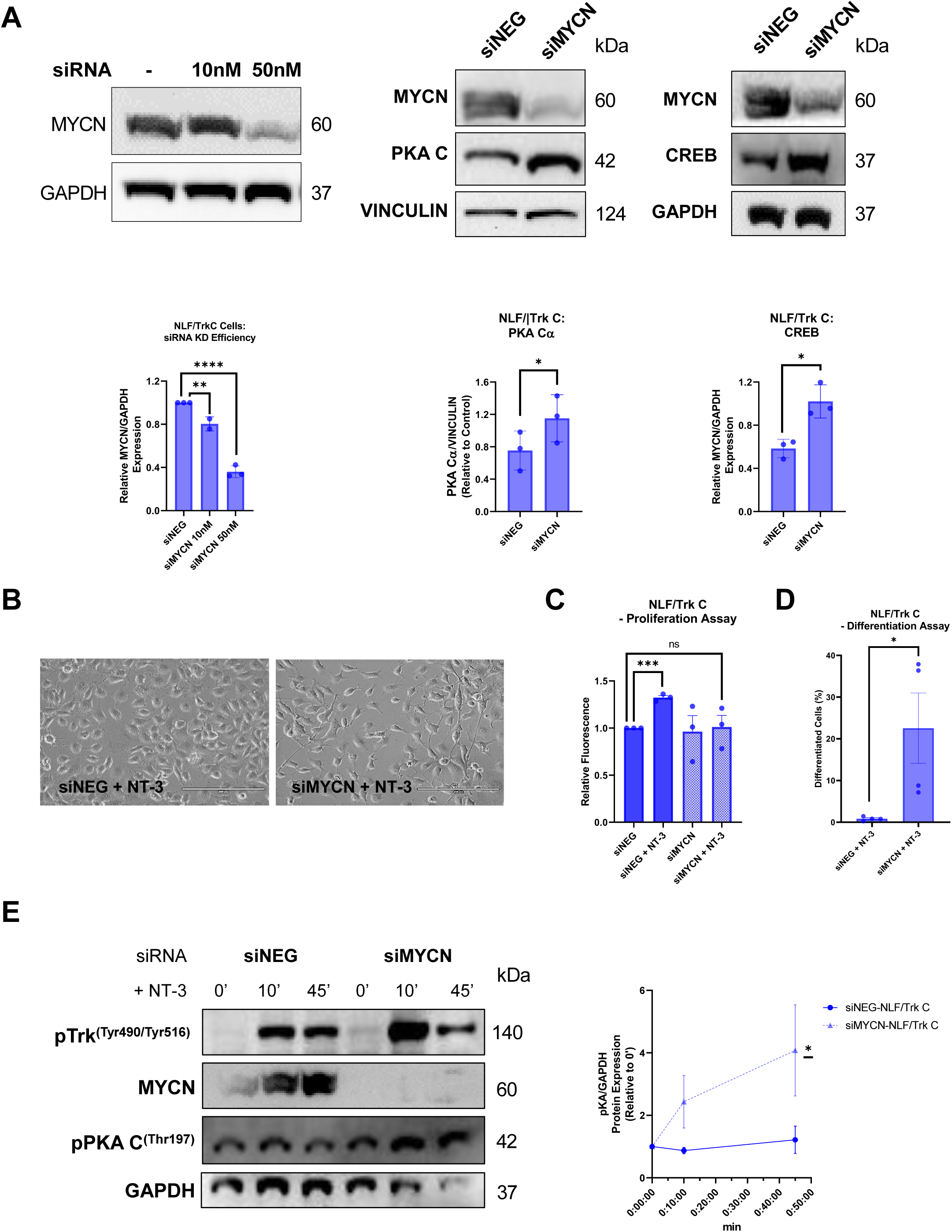
Knockdown of MYCN results in differentiation and upregulation of PKA/CREB expression in the MYCN-amplified NLF/TrkC cells. **A)** Western blot analysis of MYCN, PKA C and CREB protein expression following siRNA mediated knockdown of MYCN (siMYCN; 50 nM) or non-targeting control (siNEG) for 24 h in NLF/TrkC cells. Vinculin or GAPDH acted as loading control. Data presented as relative protein expression compared to loading control. **B)** Representative images of phenotypic observation by light microscopy after 5 days of siRNA mediated knockdown of MYCN or non-targeting control in NLF/TrkC cells. Magnification: X20; scalebar: 200 µm. **C)** Quantitation of mean fluorescence as a measure of cell proliferation at 72h post NT-3 (100 ng/ml) stimulation using CyQuant assay in NLF/TrkC cells with siRNA mediated knockdown of MYCN or non-targeting control. **D)** Quantification of neuronal differentiation with NeuronJ following NT-3 (100 ng/ml) treatment for 5 days in NLF/TrkC cells with siRNA mediated knockdown of MYCN or non-targeting control **E)** Western blot analysis of MYCN, phospho-TrkA^(Tyr490)^/TrkB^(Tyr516)^ and phospho-PKA C^(Thr197)^ in NLF/TrkC cells following 24 h of siRNA mediated knockdown of MYCN or non-targeting control treated with NT-3 (100 ng/ml) for 0-10-45 min. GAPDH acted as loading control. Data is presented as mean +/-SD of at least three independent experiments (n=3) (p < 0.05 = *, p < 0.001 = **, p < 0.0001 = ***).

Given the observation that knockdown of MYCN leads to an increase in PKA C-α and CREB expression, we further examined the impact of MYCN knockdown on PKA pathway activity. Time-course stimulation with NT-3 in NLF/TrkC cells treated with either siRNA-mediated knockdown of MYCN or a non-targeting control revealed differences in PKA activation (Fig. 4E). When MYCN was knocked down, NLF/TrkC cells exhibited increased PKA pathway activation upon NT-3 stimulation compared to the negative control. This enhanced PKA activation upon ligand stimulation, mirrors the observations in MYCN non-amplified SH-SY5Y/TrkC cells, indicating that MYCN plays a crucial role in regulating this pathway.

Signal transduction pathways mediate cellular responses and influence phenotypic changes. Phenotypic analysis of siMYCN-NLF/TrkC cells stimulated with NT-3, showed an increase in neuronal differentiation and no concomitant increase in cell number compared to siNEG-NLF/TrkC cells (Fig 4B-D).

To corroborate these findings, we also studied TrkC signalling in the MYCN-amplified IMR-32 neuroblastoma cells with high endogenous TrkC expression. Upon NT-3 treatment of IMR-32 cells, TrkC was phosphorylated and similarly, to NLF/TrkC cells they showed no significant change in phospho-PKA levels (Fig 5A). NT-3 stimulation led to an increase in cell number and no marked change in neuronal differentiation. Conversely, exogenous activation of the PKA pathway with db-cAMP reduced cell number and promoted neuronal differentiation in these cells (Fig5B-D). These results confirm that MYCN status alters NT-3/TrkC signalling and influences cell fate decisions in neuroblastoma cells.

**Figure 5.**
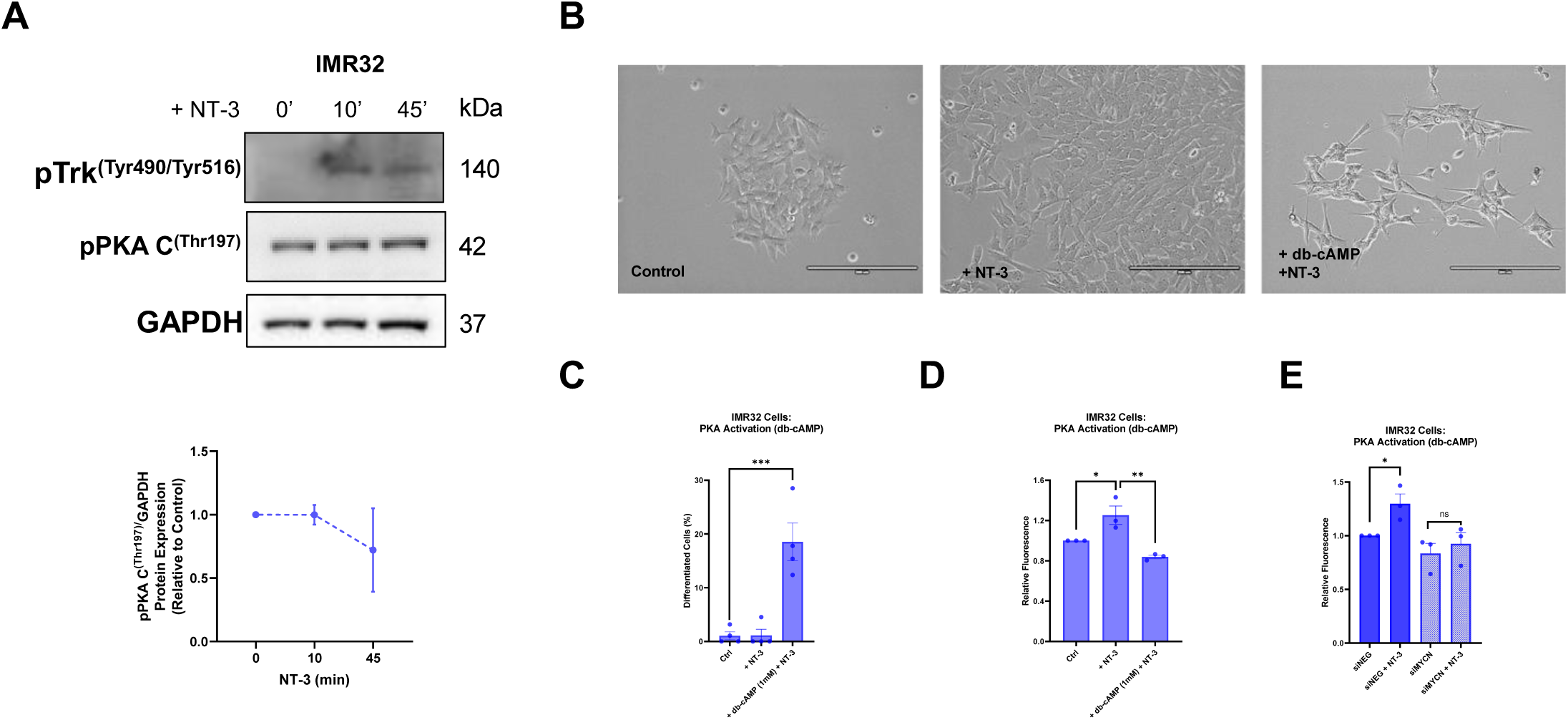
Reactivation of the PKA pathway in the MYCN-amplified IMR-32 cells with high basal TrkC expression leads to differentiation. **A)** Western blot analysis of phospho-TrkA ^(Tyr490)^/TrkB ^(Tyr516)^ and phospho-PKA C^(Thr197)^ IMR-32 cells following 0-10-45 min stimulation with NT-3 (100 ng/ml). GAPDH acted as loading control. **B)** Phenotypic observation by light microscopy after 5 days of treatment with NT-3 (100 ng/ml) and/or db-cAMP (1 mM). Magnification: X20; scalebar: 200 µm. **C)** Quantification of neuronal differentiation with NeuronJ following NT-3 (100 ng/ml) and/or db-cAMP (1 mM) treatment for 4-5 days in IMR-32 cells **D)** Quantitation of mean fluorescence as a measure of cell proliferation at 72h post NT-3 (100 ng/ml) stimulation and/or db-cAMP (1 mM) treatment using CyQuant assay (n =3) or **E)** following siRNA mediated knockdown of MYCN or non-targeting control. Data shown as mean +/-SD, (p < 0.05 = *, p < 0.001 = **, p < 0.0001 = ***).

**Figure 6.**
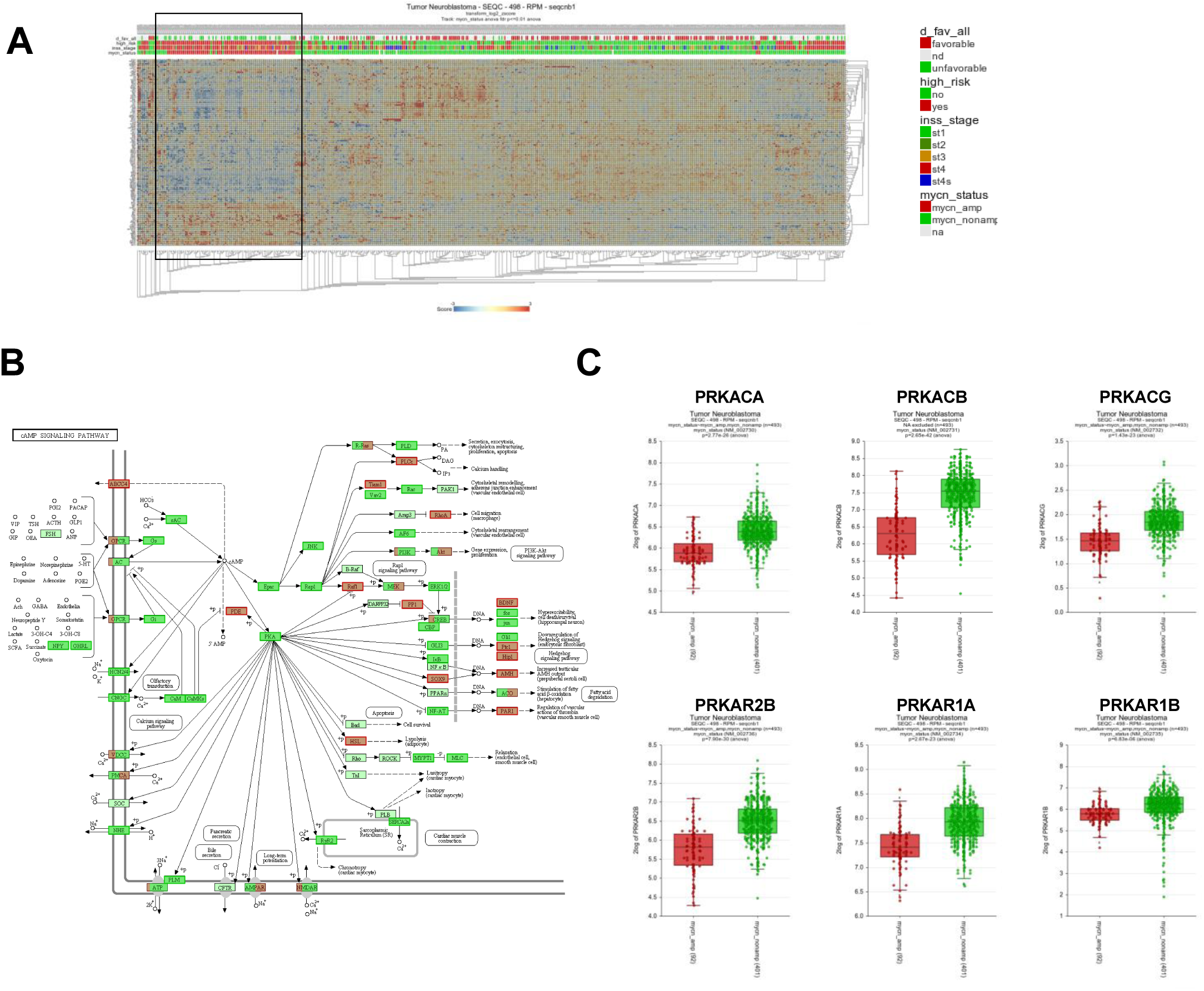
Genes related to the cAMP/PKA/CREB pathway are suppressed in MYCN-amplified patients. **A)** Heat map representing the relative expression (mRNA) of cAMP/PKA/CREB pathway related genes within a cohort of 498 neuroblastoma patients (http://r2.amc.nl; SEQC-498-RPM-seqcnb1 dataset). Each column corresponds to one patient. The cluster of MYCN amplified patients are highlighted by a square. **B)** mRNA expression of components of the cAMP/PKA/CREB pathway (KEGG: hsa04024) in the neuroblastoma patient cohort. Green colour indicates significantly higher expression of genes in the MYCN non-amplified patients, and red colour indicates significantly higher expression of genes in the MYCN-amplified patients. (Light mint green indicates no significant difference between MYCN amplified and non-amplified patients.) **C)** mRNA expression of PKA related genes in MYCN-amplified (red) vs MYCN non-amplified (green) neuroblastoma patients. Panel A - C created by using the R2: Genomics Analysis and Visualization Platform (http://r2.amc.nl).

### Neuroblastoma patients with MYCN amplification show decreased expression of PKA pathway related genes

To further examine the connection between MYCN status in neuroblastoma and cAMP/PKA/CREB pathway activity, we analysed a publicly available patient dataset comprising of 498 neuroblastoma patients (http://r2.amc.nl; SEQC-498-RPM-seqcnb1 neuroblastoma dataset). The dataset includes clinical annotations of MYCN amplification status, INSS staging, high-risk disease, age at diagnosis, disease free survival, among others (32). Of the 498 patients, 92 have MYCN amplification and 401 are MYCN non-amplified (5 not defined). Gene set analysis of the cAMP/PKA/CREB signalling pathway (hsa04024) showed that MYCN-amplified patients and the high-risk group cluster separately to MYCN non-amplified patients and have lower expression of genes involved in this pathway (Fig 7A). Mapping of gene expression data onto the cAMP signalling pathway (KEGG: hsa04024) revealed that the majority of proteins involved in this pathway are more highly expressed in MYCN non-amplified patients (Fig. 7B). Furthermore, 6 out of 7 key subunits of PKA, i.e., PKA C-α (gene: PRKACA), PKA C-β (gene: PRKACB), PKA C-γ (gene: PRKACG), PKA RI-α (gene: PRKAR1A), PKA RI-β (gene: PRKAR1B), PKA RII-β (gene: PRKAR2B), also display significantly lower gene expression in MYCN-amplified patients. There is no significant difference in the expression of the subunit PKA RII-α / gene: PRKAR2A. These findings suggest an interplay between MYCN amplification status and cAMP/PKA/CREB pathway activity in neuroblastoma.

**Figure 7.**
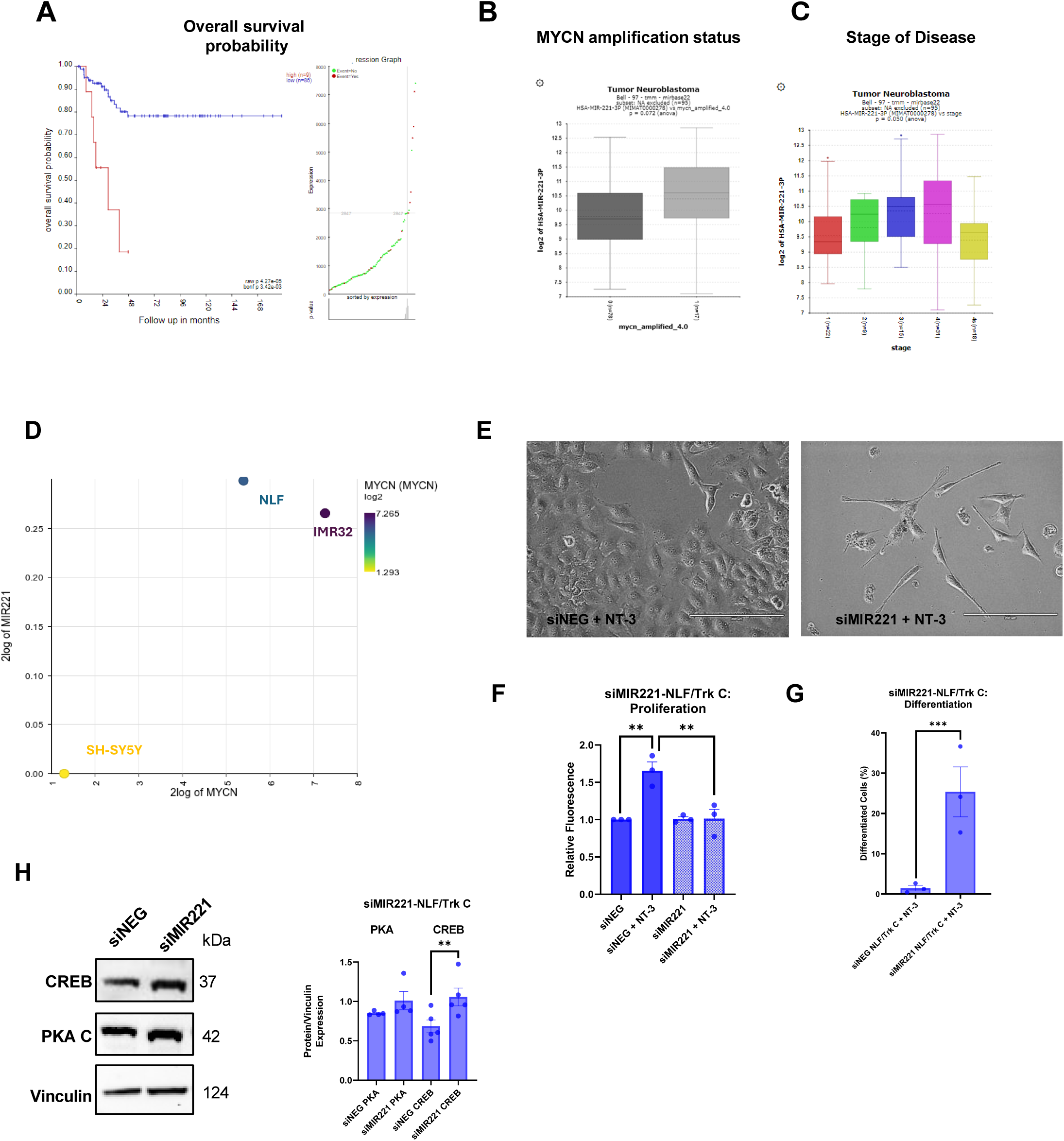
microRNA-221 positively correlates with MYCN expression in neuroblastoma and its downregulation increases CREB expression and neuronal differentiation. **A)** Kaplan-Meier curves representing the overall survival of patients with high (red) vs low (blue) mir-221 expression. (KaplanScan method was employed to differentiate high vs low expression.) **B)** Expression of mir-221 in MYCN non-amplified (red) vs MYCN-amplified (pink) neuroblastoma patients (Bell dataset) (36) **C)** Expression of mir-221 across different stages of neuroblastoma. **D)** Correlation of MYCN levels with miR221 in neuroblastoma cell lines used in this study (Maris-41-fpkm-rsg001) (41). **E)** Representative images of phenotypic observation by light microscopy after 5 days of treatment with NT-3 (100 ng/ml) in NLF/TrkC cells with siRNA mediated knockdown of MIR221 (50 nM) or non-targeting control. Magnification: X20; scalebar :200 µm. **F)** Cell proliferation measurement by CyQuant in NLF/TrkC cells following siRNA mediated knockdown of MIR221 (50 nM) or non-targeting control (siNEG) following treatment with NT-3 (100 ng/ml) for 72 h. Relative fluorescence was measured as a measure of cell number **G)** Quantification of neuronal differentiation with NeuronJ in NLF/TrkC cells with siRNA mediated knockdown of MIR221 (50 nM) or non-targeting control (siNEG) following treatment with NT-3 (100 ng/ml) for 5 days. **H)** Western blot analysis of PKA C and CREB protein expression following siRNA mediated knockdown of MIR221 (50 nM) or non-targeting control (siNEG) for 24 h in NLF/TrkC cells. Data presented as protein expression compared to loading control (vinculin). Graphs show as mean +/-SD of at least 3 independent experiments (p < 0.05 = *, p < 0.001 = **). Panel A-D created by using the R2: Genomics Analysis and Visualization Platform (http://r2.amc.nl).

### microRNA-221 positively correlates with MYCN expression in neuroblastoma and its downregulation increases PKA/CREB expression and neuronal differentiation

One mechanism by which MYCN rewires the cellular network is through the regulation of non-coding RNAs, including long non-coding RNAs, circular RNAs, and microRNAs (33–35). Numerous studies have shown that the expression of microRNAs varies depending on MYCN levels in neuroblastoma (35–38). Of particular interest, miR-221 is a microRNA, previously demonstrated to be upregulated in neuroblastoma tumours that overexpress MYCN. Studies have shown its induction by MYCN in neuroblastoma, and also, it positively correlates with poor patient survival (37,39). Additionally, its suppression is considered important for the activation of cAMP/PKA/CREB pathway and neuronal differentiation in glioma cells (40). Based on this, we investigated miR-221 as a potential mechanism of regulation.

Further bioinformatic analysis of miR-221 in patient data using the Bell dataset (36) highlighted significant differences in miR-221 expression across subtypes of neuroblastoma patients. Elevated miR-221 expression was associated with unfavourable characteristics, including MYCN amplification, decreased overall survival, and stage 4 disease (Fig. A-C).

Additionally, utilising publicly available transcriptomic data of neuroblastoma cell lines (Maris-41-fpkm-rsg001) (41), we observed a strong positive correlation between the expression levels of MYCN and miR-221 in SH-SY5Y, NLF and IMR32 neuroblastoma cells used in this study (R-value=0.916) (Fig. 7D). Knockdown of miR-221 in NLF/TrkC cells treated with NT-3 induced a more favourable phenotype in the cells by promotion of neuronal differentiation and mitigation of cell proliferation. Furthermore, the knockdown demonstrated an increase in total CREB expression (PKA levels were not significantly different). Collectively, these findings suggest that high levels of miR-221 in MYCN amplified neuroblastoma cells suppress the PKA/CREB pathway, consequentially inhibiting TrkC-mediated neuronal differentiation (Fig. 8).

**Figure 8.**
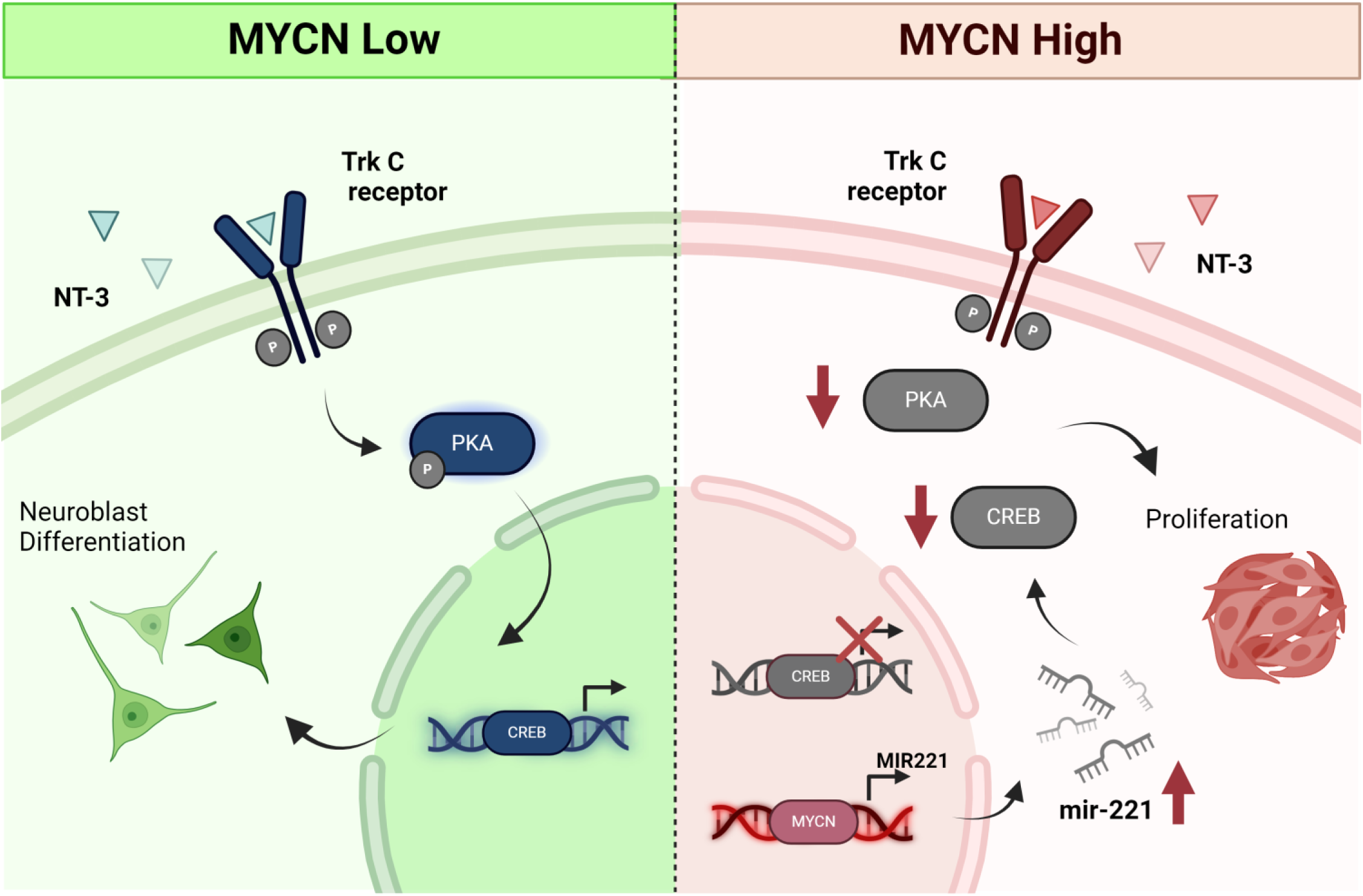
TrkC mediated signalling in NB. PKA signalling is crucial for inducing TrkC-mediated differentiation in non-MYCN-amplified NB cells. When MYCN is overexpressed, MYCN inhibits the PKA/CREB signalling pathway via miR-221.

## Discussion

This study provides a comprehensive evaluation of NT-3/TrkC signalling in the context of neuroblastoma. Early studies on TrkC in neuroblastoma highlighted TrkC expression to be more prevalent in Stage 1, 2, and 4S tumours (37%), in comparison to only 12% in stage 3 and 4 tumours. Additionally, Ryden et al. showed that TrkC expression is seen with lower age and lower stage of disease, both suggesting its expression to be associated with a more favourable prognosis (16,42). Taking into consideration TrkC’s ligand, NT-3, Bouzas-Rodriguez, showed that NT-3 is upregulated in a large fraction of aggressive human neuroblastomas, and additional studies highlight NT-3’s clinical significance to be different with regard to TrkC expression and MYCN amplification, thereby, suggesting it’s activity to be dependent on the molecular environment (17,43). Likewise, TrkC has also shown to have opposing effects in other cancers, displaying both oncogenic and tumour suppressive effects, emphasising its context dependent behaviour (18,19,21,22).

Moreover, the above-mentioned studies, largely focus on the mRNA expression of TrkC and NT-3 in neuroblastoma. Little is known about downstream signalling events following TrkC receptor activation in neuroblastoma cells, beyond putative activation of ERK, AKT, PLCy. Utilising a quantitative mass spectrometry-based phosphoproteomic approach, our study provides a comprehensive profile of NT-3/TrkC signalling in neuroblastoma and its impact on cellular phenotype. Additionally, the use of temporal profiling provide insight into early-intermediate- and late signalling events downstream of receptor activation, offering a well characterised dataset of the signalling network.

MYCN is a crucial consideration when studying neuroblastoma. The amplification of this oncogene is seen in half of high-risk cases and serves as the most significant predictor of poor prognosis, guiding patient stratification in clinical practice (24,44–46). At the cellular level, MYCN amplification has been shown to exert profound reprogramming of cellular signalling networks, metabolic rewiring, immune surveillance functions, epigenetics processes, gene expression and most recently repression of circular RNAs (34,47–50). Duffy et al, showed MYCN as a global suppressor of cellular signalling in neuroblastoma and distinct differences in the cellular network between endogenous MYCN and even between overexpressed MYCN and amplified MYCN (38) .

With respect to this, our study takes into consideration different MYCN levels in neuroblastoma. Through the generation of TrkC overexpressing neuroblastoma cell lines, that demonstrate varying levels of MYCN expression, this work permits the molecular and phenotypic characterisation of TrkC in different neuroblastoma cellular contexts and allows a specific focus on understanding how MYCN rewires the TrkC signalling network. Indeed, we also observed a MYCN-dependent influence on downstream signalling and cellular phenotype. Cell lines with MYCN overexpression or amplification showed a proliferative phenotype when TrkC pathway was activated compared to differentiation in MYCN non-amplified neuroblastoma cells. This phenotype was interconverted upon siRNA mediated KD of MYCN. In concordance with Duffy et al., MYCN also altered downstream cellular signalling. This was observed on a global level, where PCA clustering of NT-3/TrkC phosphoproteomics data showed distinct separation based on cell lines with different MYCN status.

In particular, our study identified differential activation of PKA/CREB pathways in response to NT-3 stimulation and highlighted the significance of this pathway in neuroblastoma cell fate determination. Previous research also highlights this pathways involvement in neuronal differentiation of neural stem cells, hippocampal progenitor cells and also in promoting the differentiation of pluripotent stem cells (51–54). Moreover, Sánchez et al. also recognised cAMP mediated neurite formation in neuroblastoma cells. Their results demonstrate both PKA and PI3K activation to be essential for this phenotype (55). PI3K/AKT related pathways were also enriched in KSEA of SH-SY5Y/TrkC cells in our data. However, when investigated further, we noticed no significant difference in PI3K/AKT pathway activity between SH-SY5Y/TrkC and NLF/TrkC cells following NT-3 stimulation. Also, pharmacological inhibition of PI3K/AKT or constitutive activation of AKT did not alter neuronal differentiation in our cells.

siRNA mediated KD of MYCN in MYCN-amplified cells resulted in an upregulation of CREB protein expression suggesting MYCN status to alter PKA/CREB signalling and consequently NT-3/TrkC mediated cell fate decisions. Furthermore, our findings were supported by patient data, where MYCN amplified neuroblastoma patients demonstrated lower mRNA expression of PKA related genes and overall suppression of components of the cAMP/PKA/CREB signalling pathway. The correlation of our in vitro findings with clinical data provides assurance that our results accurately reflect the role of MYCN biology within the clinical context of neuroblastoma. This finding is also supported by other studies that highlight CREB’s involvement in neuronal differentiation and suppresses neuroblastoma tumour growth (56,57).

One way in which MYCN has been shown to rewire the cellular network is via regulation of noncoding RNAs such as long noncoding RNAs, circular RNAs and microRNAs(33–35). Many studies highlight microRNAs to be differently expressed dependent on MYCN levels in neuroblastoma(35–38). This study suggests MYCN to regulate the cAMP/PKA/CREB pathway and suppress neuronal differentiation via upregulation of microRNA 221. MicroRNA 221 has been previously observed to be upregulated by MYCN in neuroblastoma, correlating with unfavourable patient outcomes (36,37,39). Notably, its downregulation has also been shown to activate the cAMP/PKA/CREB pathway and promote neuronal differentiation in glioma cells (40). Additionally, miR-221 may also suppresses NT-3/TrkC mediated differentiation in neuroblastoma patients with high MYCN via regulation of NT-3. Investigation into potential targets of miR-221 using the TargetScan database revealed NT-3 to also be a target of this microRNA (58). Further exploration into this regulatory axis may prove informative for understanding difference in NT-3/TrkC mediated differentiation in neuroblastoma.

Importantly, this study also identifies a novel therapeutic approach in neuroblastoma. Differentiation of neuroblasts to mature benign neuronal cells is a therapeutic aim in the treatment of neuroblastoma. Low stage tumours show a more differentiated phenotype, and it is also proposed as a mechanism driving the spontaneous regression of Stage 4S tumours (59). Differentiation is currently achieved in the clinic during maintenance therapy by administration of all-trans retinoic acid to patients (60,61). Owing to, its effectiveness being limited to minimal residual disease much research efforts are ongoing to identify novel differentiation agents or agents to potentiate the effects of retinoic acid. Exogenous activation of the cAMP/PKA/CREB pathway using cAMP analogues may prove therapeutically beneficial in this regard. In fact, cAMP agonist, tocladesine (8-Cl-cAMP), has been previously used in stage I/II clinical trials in colorectal cancer, multiple myeloma, and plasma cell neoplasm (NCT00004902, NCT00021268), highlighting its clinical translatability. Further exploration is warranted to fully elucidate its therapeutic potential in differentiation of neuroblastoma.

In conclusion, this research provides a comprehensive analysis of the NT-3/TrkC signalling network in neuroblastoma and how MYCN can alter the signalling network and cell fate decisions in this context. We identified the cAMP/PKA/CREB pathway to be important for TrkC mediated neuronal differentiation of neuroblastoma cells and suggest MYCN to be suppressing this pathway via upregulation of microRNA 221 (Figure 8). Manipulation of this pathway has potential for therapeutic benefit in differentiation treatment of MYCN-amplified neuroblastomas.

## Materials and Methods

### Antibodies, ligands and inhibitors

Antibodies used include TrkC (C44H5) (#3376), Phospho-TrkA ^(Tyr490)^/TrkB ^(Tyr516)^ (C35G9) (#4619), Phospho-ERK (^Thr202/Tyr204)^ (#4370), Phospho-Akt ^(Ser473)^ (#4060), Phospho-PKA C^(Thr197)^ (#4781) Vinculin (#4650), PKA C-α (#4782), CREB (#9197), and GAPDH (14C10) (#2118), from Cell Signalling Technology. V5-tag monoclonal antibody (#R960-25) and Phospho-PLCγ^(Tyr783)^ (#44-696) were from Invitrogen and the MYCN antibody (#53993) used is from Santa Cruz Biotechnology. Secondary horseradish peroxidase–conjugated antibodies against rabbit (#7074) or mouse (#7076) immunoglobulin G (IgG) were from Cell Signalling Technology.

Recombinant Human NT-3 (#450-03) was obtained from Peprotech. The PKA inhibitor, H 89 2HCl (#S1582) and the PKA activator, dibutyryl-cAMP (#S7858) were purchased from Selleck Chemicals.

### Cell culture

The human neuroblastoma cell lines SH-SY5Y, NBLS and IMR-32 were obtained as a gift from Frank Westermann (Deutsches Krebsforschungszentrum (DKFZ), Heidelberg, DE) and the NLF cell line was obtained from Kerafast (ECP008). Cell lines were cultured in RPMI 1640 (Gibco) supplemented with 10% (v/v) foetal bovine serum (Gibco), 2 mM L-glutamine (Gibco), and penicillin (100 U/ml) and streptomycin (100 μg/ml) (Gibco). The media for cell lines expressing the pLX302/NTRK3 plasmid were additionally supplemented with 1 μg/ml puromycin (Sigma-Aldrich) to maintain plasmid expression. All cell lines were routinely tested for mycoplasma.

### Generation of pLX302/NTRK3 cell lines

Cells were transfected with 5 μg of DNA using jetPrime transfection reagent (Polyplus) according to the manufacturer’s instructions. After 24 h, the media was changed to RPMI 1640 full media containing puromycin (1 μg/ml) to allow antibiotic selection of positively transfected cells.

### Cell stimulation and lysis

Cells were serum starved with media containing 0.1% FBS for 6 h prior to treatment. After starvation, the cells were treated with NT-3 (100 ng/ml) +/-inhibitor for time-course stimulation. Cells were lysed in lysis buffer (1% Triton x100, 20 mM Tris-HCl pH 7.5, 150 mM NaCl, 1 mM MgCl2), supplemented with protease inhibitor Complete Mini (Roche) and phosphatase inhibitor PhosStop (Roche). Lysates were cleared by centrifugation (14,000 rpm, 10 min, 4°C) and the supernatants were stored at -20°C until analysis.

### Western blotting

SDS-polyacrylamide gel electrophoresis (SDS-PAGE) and Western blotting were performed using the Bolt Mini Gel Tank and Bolt Bis-Tris Precast 10% gels (Invitrogen). Gels were transferred onto PVDF membranes using the XCell II Blot Module system (Invitrogen). Membranes were blocked in 5% non-fat dried milk (Sigma-Aldrich) for 1 h at room temperature prior to overnight incubation at 4℃ with primary antibody diluted in bovine serum albumin (BSA) (1:1000). Membranes were washed in TBS-Tween (3 x 5 min) then incubated with the corresponding secondary antibodies diluted in 5% milk (1:5000) for 1 h at room temperature. Blots were washed and then developed using the iBright imaging system (ThermoFisher Scientific) with ECL or SuperSignal West Femto Chemiluminescent Substrate (ThermoFisher Scientific). Quantification of blots was achieved using ImageJ software v1.44p (http://imagej.nih.gov/ij).

### Cell proliferation

Cells were seeded in technical triplicates at 5,000 cells/well in 96-well plate format. After 4 h, cells were treated with NT-3 (100ng/ml) +/-inhibitors. At 72 h post treatment, cell proliferation was measured using the CyQUANT™ Assay (ThermoFisher Scientific) according to the manufacturer instructions. Fluorescence was measured using the Spetramax M3 Plate reader.

### Neurite outgrowth analysis

Cells (5 x 10^3^ /ml) were treated with NT-3 (100 ng/ml) +/-inhibitor in 6-well plate format. Media and treatment were refreshed every 2 days. Using light microscopy, images were taken at day 5 using EVOS FL Imaging System microscope (20X magnification). Neurite elongation and cell number was measured on three images from each experimental condition using NeuronJ Software (62)(63) and cellpose3 package in Python (64), respectively. NBLS/Trk C cells were counted manually due to segmentation issues in cellpose3.

### Luciferase reporter assay

The CRE/CREB Reporter Kit (cAMP/PKA Signalling Pathway) (Generon) and BPS Two-Step Luciferase (Firefly & Renilla) Assay System was used according to manufacturer instructions and measured using the Spetramax M3 Plate reader.

### siRNA-mediated knockdown

The MYCN (#L-003913-01-0005), MIR221 Silencer^®^ Select (#4390771) and negative control (D-001810-10-05) ON-TARGETplus SMARTpool siRNAs were purchased from ThermoFisher Scientific. Transfection was performed using jetPRIME reagent (Polyplus) according to manufacturer instructions.

### Liquid chromatography tandem mass spectrometry (LC-MS/MS)

Samples for mass spectrometry were prepared as described in Maher et al (65). Samples were run on a Bruker timsTof Pro mass spectrometer connected to a Evosep One liquid chromatography system as described previously (62)

The mass spectrometer raw files were searched against the Homo sapiens subset of the Uniprot Swissprot database (reviewed) using the search engine FragPipe (Version 18) (62).

### Bioinformatics

The mass spectrometry proteomics data have been deposited to the ProteomeXchange Consortium via the PRIDE(66) partner repository with the dataset identifier PXD054441. Data analysis was performed in R (Version 4.1.2). LFQ intensities were log2-transformed. Proteins/phosphosites with more that 80% missing values in all conditions were filtered out. Missing values were imputed using the group mean imputation with normal distribution correction and tail-based imputation approach. Analysis of differently expressed phosphosites and proteins was performed using the limma package in R/Bioconductor (67) with adjusted p.value < 0.05 and absolute fold change > 1.5 as the cutoffs for a phosphosite to be considered significantly different compared to the zero timepoint of each experimental condition. Kinase substrate enrichment analysis was performed using the KSEAapp R package (68,69) with default parameters.

### Patient data

Analysis of neuroblastoma patient gene expression data was performed using the R2 Genomics Analysis and Visualization Platform (http://r2.amc.nl). The SEQC-498-RPM-seqcnb1, Kocak - 649 - custom - ag44kcwolf data set and TARGET - 161 – fpkm bulk RNA-Seq datasets were used. Gene set analysis of KEGG pathways (hsa04024) was performed (p-value cut-off < 0.05). Analysis of individual genes was done by comparing the log2 gene expression separated by MYCN-status (MYCN amplified vs MYCN non amplified, NA samples were excluded). Expression was considered significantly different if p-value < 0.05.

### Statistical analysis

All experiments were performed in three biological replicates. Statistical analysis for cell-based assays and Western blots were conducted using GraphPad Prism (version 8.0.2). Data is expressed as mean ± SD. Student t-test was used for comparison between two groups or One-way Anova for more than two groups. p-value < 0.05 were considered statistically significant. *p-value <0.05, **p-value < 0.01, ***p-value < 0.001.

## Acknowledgements

This research was funded by the Irish Research Council under grant number GOIPG/2020/1361; the Comprehensive Molecular Analytical Platform (CMAP) under the Science Foundation Ireland Research Infrastructure Programme (reference 18/RI/5702); the Precision Oncology Ireland grant 18/SPP/3522 by Science Foundation Ireland and Children’s Health Ireland; and with the financial support of Children’s Health Foundation and under the management of Science Foundation Ireland under the Frontiers for the Future Programme Grant Number 21/FFP-P/10130.

## Author contributions

M.H. conceived the study and supervised the experiments, S.M. conducted the experiments. K.W performed the mass spectrometry. S.M., M.H, K.W and V.Z. analysed the results. S.M and M.H wrote the manuscript. All authors reviewed and approved the final version of the manuscript.

## Data Availability

The mass spectrometry proteomics data have been deposited to the ProteomeXchange Consortium via the PRIDE [1] partner repository with the dataset identifier PXD054441.

